# Analog of the Hutchinson equation in biophysical neurodynamics: from the Morris-Lecar model to a delay differential equation

**DOI:** 10.1101/2022.01.15.476459

**Authors:** Alexander Paraskevov

**Affiliations:** Institute for Information Transmission Problems, 127051 Moscow, Russia

**Keywords:** Morris-Lecar model, Neuronal excitability, Constant current stimulation, Periodic spiking, Time-delay system, Delay differential equation

## Abstract

Starting with the classical biophysical Morris-Lecar model of neuronal excitability, we introduce a functional analog of the Hutchinson equation initially obtained for population dynamics with delayed negative feedback. It is shown that the resulting equation with a fixed time delay qualitatively reproduces the dynamics of the original model upon direct current stimulation, preserving both the initial type of neuronal excitability and biophysically realistic spike shape within a wide range of the delay values. If the delay becomes very small (2 ms or less), the simplified delay-based model exhibits a distinct transition from the 1st to the 2nd excitability type.

## 1. Introduction

This work addresses simplification in describing nonlinear neuronal dynamics by functional analogy with a well-known equation for population dynamics. Guided by the principle that science is a search for the simple in the complex, we have taken a classical biophysical neuron model and simplified it substantially while preserving its essential properties.

A bit of history to start. In 1948, ecologist G.E. Hutchinson, studying dynamics of a population size, generalized the Verhulst logistic equation

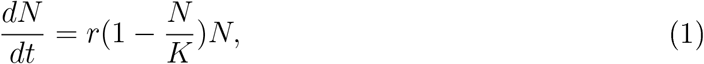

where *N* is the population size, *r* is the population growth rate, and *K* is the average (and asymptotic) population size. Provided with initial condition *N* (*t* = *t*_0_) = *N*_0_, the equation has a well-known exact solution in the form of logistic function,

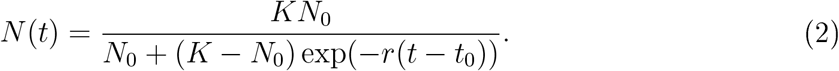

To account for the period of sexual maturation of individuals, Hutchinson added a constant time delay *t*_*del*_ to the multiplier limiting population growth [1],

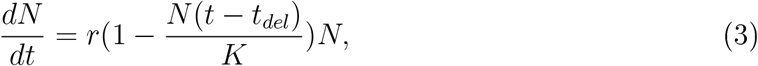

that led to the emergence of nontrivial self-oscillatory solutions (Fig. 1, left graph) [2]. In particular, it turned out that the period of such self-oscillations can be much longer than *t*_*del*_. By that time, similar self-oscillatory solutions were already known in the Lotka-Volterra model describing the dynamics of two populations (conventionally, “predators” and “preys”) with a one-sided antagonistic interaction between them. However, the emergence of such self-oscillations for a single delay-differential equation [3] came as a surprise, so the denominative “Hutchinson equation” arose.

**Figure 1.**
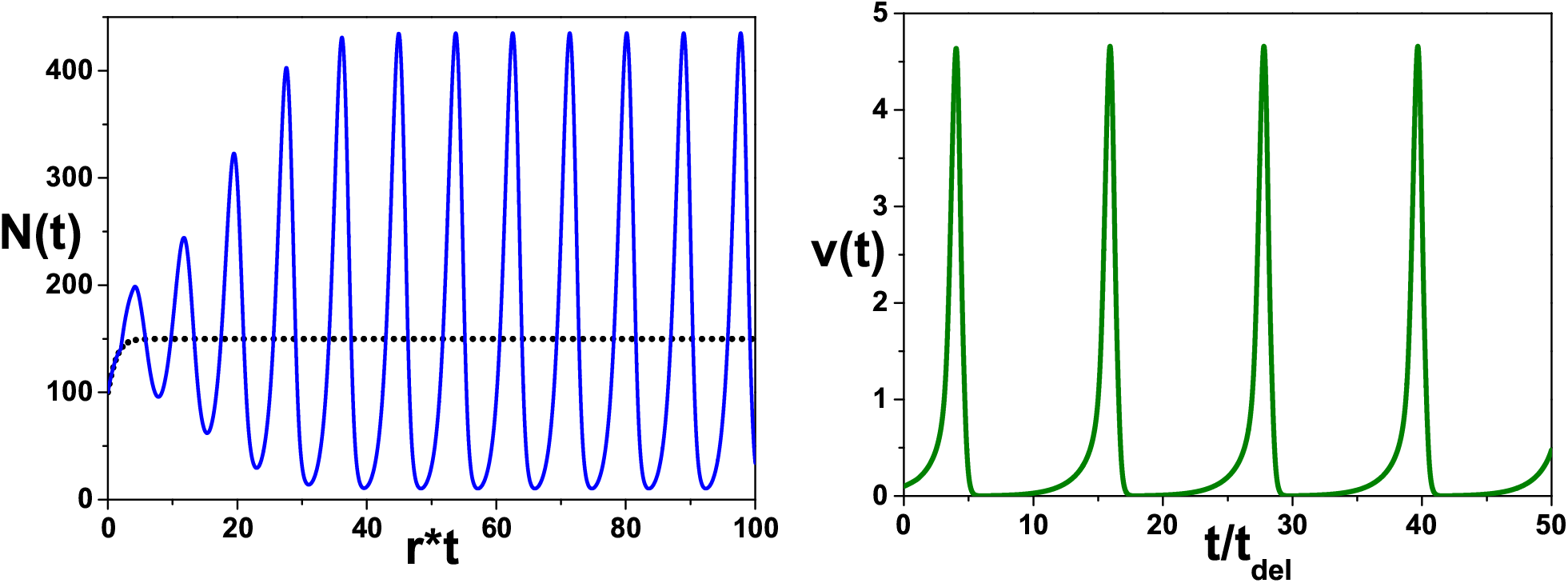
Left graph: An example of self-oscillatory solution (blue solid line) of the Hutchinson equation (3) for *N*_0_ = 100, *K* = 150, *r* = 0.4, and *t*_*del*_ = 5. The black dotted line is the corresponding solution (2) of the Verhulst logistic equation (1) with *t*_0_ = 0. Note that at *t ≤ t*_*del*_ both solutions coincide. Right graph: An example of self-oscillatory solution for the Mayorov-Myshkin model (5), where *λ* = 3, *R*_1_ = 1, *R*_2_ = 2.2.

In the same 1948, A.L. Hodgkin, as a result of classical experiments on stimulating isolated axons with direct current, proposed to classify spiking excitability of an axon into the following three types or classes [4]. Type 1: the average frequency *f* (*I*_*stim*_) of spike generation, as a function of constant stimulating current *I*_*stim*_, can be arbitrarily small (or, in other words, it starts growing from zero continuously). Type 2: function *f* (*I*_*stim*_) is discontinuous, i.e., it abruptly takes a non-zero minimal value. Type 3: the axon is unable to periodically generate spikes, regardless of the stimulating current value. This classification has been later transferred without changes to the entire neuron and is now generally accepted.

In this article, we show that it is possible to get a functional analog of the Hutchinson equation based on the classical biophysical Morris-Lecar (ML) model of neuronal excitability [5, 6], which takes into account the dynamics of voltage-dependent ion channels and plausibly describes the action potential (i.e., spike) waveform. In general, the spike generation is a manifestation of nonlinear impulse relaxation of the neuron transmembrane potential to a stationary physiological value, the so-called resting potential. In standard electrophysiological studies, a basic method (“protocol”) for studying the spiking response of a neuron and identifying the type of neuronal excitability is micro-electrode stimulation of the neuron by constant depolarizing current. In what follows, we imply this protocol everywhere. Mathematically, the ML model consists of a system of two nonlinear differential equations: a dynamic equation for the transmembrane potential *V* of the neuron and a relaxation equation for the dimensionless conductance *w* of potassium ions, which can also be considered as the probability of opening potassium ion channels in the neuron membrane. Spikes represent characteristic pulses of the potential *V* (see Fig. 2). The change rate of *V* depends on the current value of *w* so that the dynamics of *w* provides a negative feedback with respect to the dynamics of *V*. In turn, change rate of *w* is proportional to the difference between the current value of *w* and the “stationary” (or asymptotic) value *w*_*∞*_(*V*), to which *w* tends when the stimulation is turned off and which depends nonlinearly on *V*. It is also worth noting that, as the ML model has been originally developed for the excitability of a muscle fiber, it considers the transmembrane dynamics of calcium but not sodium ions.

**Figure 2.**
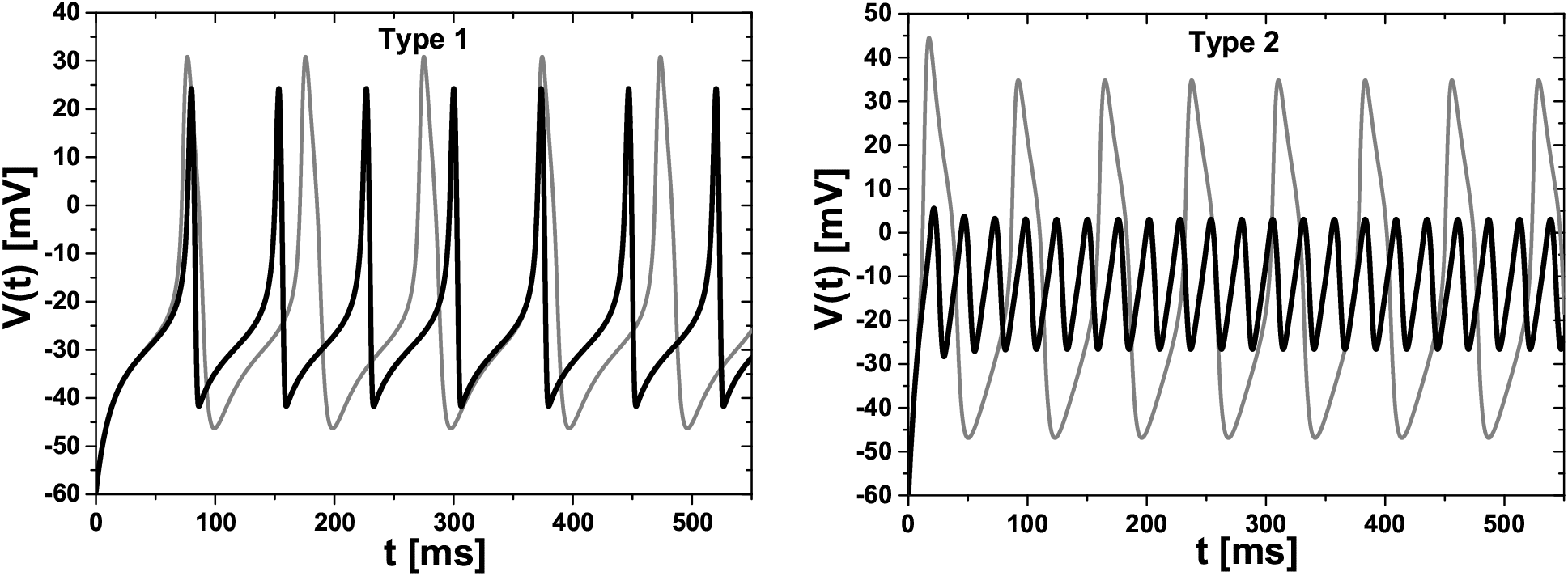
Left graph: The gray line is a numerical solution for the dynamics of neuron potential *V* (*t*) in the standard Morris-Lecar model (6) with the 1st excitability type at *I*_*stim*_ = 45 *μ*A/cm^2^. The black line is the corresponding numerical solution of the Simplified Delay-based ML (SDML) model (11) at *t*_*del*_ = 3 ms. Note that the spike shape is well preserved. Right graph: The analogous solutions for the 2nd excitability type at *I*_*stim*_ = 122 *μ*A/cm^2^ and *t*_*del*_ = 3 ms. For the 2nd type, the spike amplitude in the SDML model grows substantially with increasing *t*_*del*_ (see the bottom right graph in Fig. 3).

If one completely neglects the relaxation dynamics of *w* and equates *w*(*t*) = *w*_*∞*_(*V* (*t*)), then spike generation in the ML model does not occur. To simplify the relaxation dynamics, we have replaced the original equation for *w* by the equation *w*(*t*) = *w*_*∞*_(*V* (*t* − *t*_*del*_)), where *t*_*del*_ is a fixed time delay of several milliseconds. The ML model modified in such a way can be called as Simplified Delay-based ML (SDML) model. In this case, when the neuron is stimulated by a direct current, spikes can occur (see Fig. 2), and, importantly, the original type of neuronal excitability is preserved within a wide range of *t*_*del*_ values. Thus, it turns out to be possible formulating a biophysically realistic (with respect to the spike shape) model of neuronal excitability, consisting of only one dynamic equation for the neuron potential. This result virtually refutes a widespread belief that the minimal number of equations in a dynamic model of neuronal excitability to describe more-or-less realistic spike shape should be equal to two - as in the original ML model or in the FitzHugh-Nagumo model [7, 8], which has originated as a result of attempts to simplify the Hodgkin-Huxley model [9].

It should be noted that using a delay differential equation [3] for the description of neuronal dynamics has been already considered in the so-called Mayorov-Myshkin (MM) model [10] (see also [11, 12]),

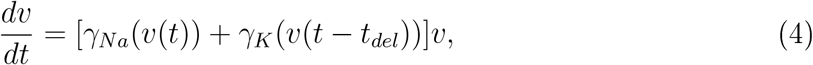

where *v* ≥ 0 is the dimensionless deviation of the membrane potential from the level of the highest membrane polarization (i.e., when the neuron is in the hyperpolarization state), and *γ*_*Na*_(*v*) = *a* − *f*_*Na*_(*v*) and *γ*_*K*_(*v*) = *f*_*K*_(*v*) − *b* are respectively the sodium and potassium conductances divided by the electrical capacitance of the membrane. Further, *f*_*Na*_(*v*) and *f*_*K*_(*v*) are positive and monotonically decreasing functions, and parameters *b* > *a* > 0 are such that *f*_*Na*_(0) *> a, f*_*K*_(0) *> b*, and *f*_*K*_(0) − *f*_*Na*_(0) *> b* − *a*. Taking *f*_*Na*_(*v*) and *f*_*K*_(*v*) in the form of the Gaussian functions, *f*_*Na*_(*v*), *f*_*K*_(*v*) *∝* exp(−*v*^2^), and taking the value of time delay *t*_*del*_ as the normalizing time scale, the authors arrived at the final equation of their model:

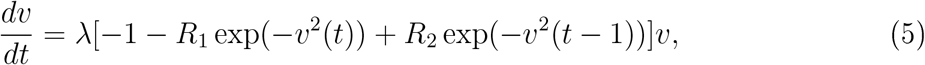

where *λ* = (*b* − *a*)*t*_*del*_, *R*_1_, and *R*_2_ are constant parameters. Their typical values are *λ* = 3, *R*_1_ = 1, and *R*_2_ = 2.2 [10]. The model plausibly describes the spike waveform, and exhibits a rich dynamic behavior including pacemaker (i.e., self-oscillation) regime (see the right graph in Fig. 1). Despite all other assumptions, the key point of the MM model is a finite delay in the dynamics of potassium conductance. This point is the same as in our simplified version of the ML model. Nevertheless, the SDML model seems a way more grounded than the MM model, which is derived from purely speculative consideration. Indeed, the parameters of the former model are adopted from the ML model and have a clear biophysical meaning, while the ones for the MM model are relatively abstract. In addition, there are many numerical studies of neuronal networks with the ML model as the single neuron model (e.g., [13–15]). In most of such studies the ML model can be readily transformed into the SDML model, but not into the MM model. This is especially true for the need to adapt the constituent model of “chemical” synaptic interaction between neurons: unlike the MM model, SDML neurons are coupled in exactly the same way (i.e., using the same synapse model) as ML neurons. Therefore, the preference for the SDML model is apparent, even though the MM model might also be quite fruitful potentially.

Finally, in a recent study [16] the standard ML neuron model has been supplemented with a synapse-autapse [17, 18], when the outgoing synapse of a neuron is simultaneously the incoming one to the same neuron, i.e., the neuron is connected through the synapse to itself. Due to the finite time of both the axonal and chemical synaptic transmissions, the synaptic current provides a delayed feedback for the neuron potential dynamics (cf. [19, 20]). The delayed feedback is functionally similar to the one we consider, though it has no relation to the conductance of potassium ions.

## 2. Standard Morris-Lecar model and its delay-based simplification

In the standard two-dimensional ML model [5, 6] (cf. [21]), the equations for dynamics of the neuronal potential *V* and for relaxation dynamics of the normalized conductance *w* of potassium ions are given by

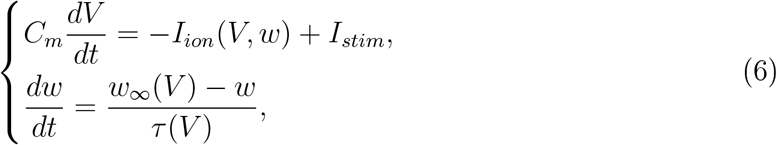

where *I*_*ion*_(*V, w*) is a sum of two ion currents and the leakage current,

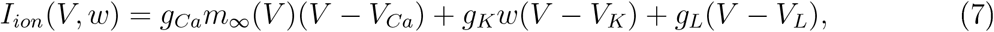

*I*_*stim*_ is an external stimulating current, and the constituent functions

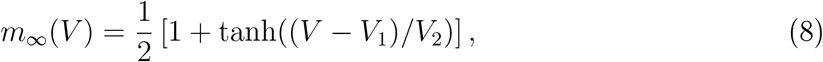

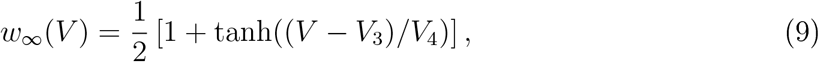

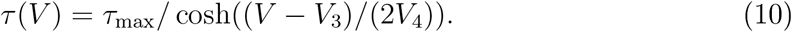

In numerical simulations reported further, we have used the following values of parameters. For the ML model of the 1st neuronal excitability type [6]: *C*_*m*_ = 20 *μ*F/cm^2^, *g*_*Ca*_ = 4 mS/cm^2^, *g*_*K*_ = 8 mS/cm^2^, *g*_*L*_ = 2 mS/cm^2^, *V*_*Ca*_ = 120 mV, *V*_*K*_ = −84 mV, *V*_*L*_ = −60 mV, *V*_1_ = −1.2 mV, *V*_2_ = 18 mV, *V*_3_ = 12 mV, *V*_4_ = 17.4 mV, *τ*_max_ = 14.925 ms. These parameters result in the resting potential value *V*_*rest*_ = −59.47 mV, which is the solution of equation *I*_*ion*_(*V, w*_*∞*_(*V*)) = 0 and is very close to *V*_*L*_ value. For the ML model of the 2nd excitability type [6]: *g*_*Ca*_ = 4.4 mS/cm^2^, *V*_3_ = 2 mV, *V*_4_ = 30 mV, *τ*_max_ = 25 ms, and all the rest parameters are the same as those for the 1st type. In turn, these parameters result in *V*_*rest*_ = −60.85 mV.

The initial conditions for all numerical simulations of the ML model in this article were as follows: *V* (*t* = 0) = *V*_*rest*_, *w*(*t* = 0) = *w*_*∞*_(*V*_*rest*_).

As described above, the ML equations (6) can be simplified to a single delay differential equation

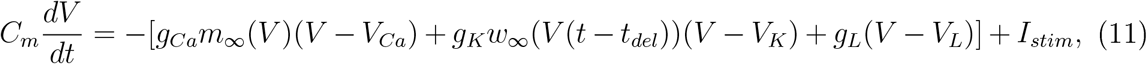

where *t*_*del*_ *>* 0 is a fixed time delay of several milliseconds. In particular, we have used *t*_*del*_ value equal 3, 6, and 9 ms. For the interval 0 *≤ t < t*_*del*_, in Eq. (11) we use *w*_*∞*_(*V*(*t*)) instead of *w*_*∞*_(*V* (*t* − *t*_*del*_)), i.e., while the time is small enough there is no delay. We refer to the model described by Eq. (11) as the Simplified Delay-based ML SDML model.

## 3. Results

We have numerically studied excitability properties of the SDML model (11) for the case of stimulating the model neuron by constant depolarizing current.

The main results are shown in Fig. 3. In particular, according to the hallmark dependence of spike generation frequency on the strength of the constant stimulating current, for the SDML model with parameters corresponding to the 1st and 2nd types of excitability for the ML model, the excitability types are preserved within the time delay interval 2.5 ms ≤ *t*_*del*_ ≤ 1 ms (blue curves on the upper graphs in Fig. 3). Defined as difference *V*_*max*_ −*V*_*min*_, where *V*_*max*_ and *V*_*min*_ are the maximal and minimal values of the neuron s potential at given stimulating current *I*_*stim*_, the spike amplitude is shown ibid, by the corresponding green curves for three different values of *t*_*del*_. In the above *t*_*del*_ interval, periodic spike generation depending on the value of *I*_*stim*_ ≥ *I*_*min*_ always starts with a discontinuous occurrence of the spike amplitude at *I*_*stim*_ = *I*_*min*_, regardless of the excitability type. Undamped periodic spiking stops at *I*_*stim*_ ≥ *I*_*max*_ [22] so that it is possible only in a finite range of stimulating current values, *I*_*min*_ ≤ *I*_*stim*_ *< I*_*max*_. In turn, upon stimulation with the minimal current necessary for periodic spike generation, *I*_*stim*_ = *I*_*min*_, the spike amplitude monotonically increases as the function of *t*_*del*_ over the entire interval 2.5 ms ≤ *t*_*del*_ ≤ 15 ms for the 1st excitability type. For the 2nd type, this dependence is not monotonic in the range of 2.5 ms ≤ *t*_*del*_ ≤ 7 ms, but it returns to monotonous growth further (see the green curves in the lower graphs in Fig. 3).

**Figure 3.**
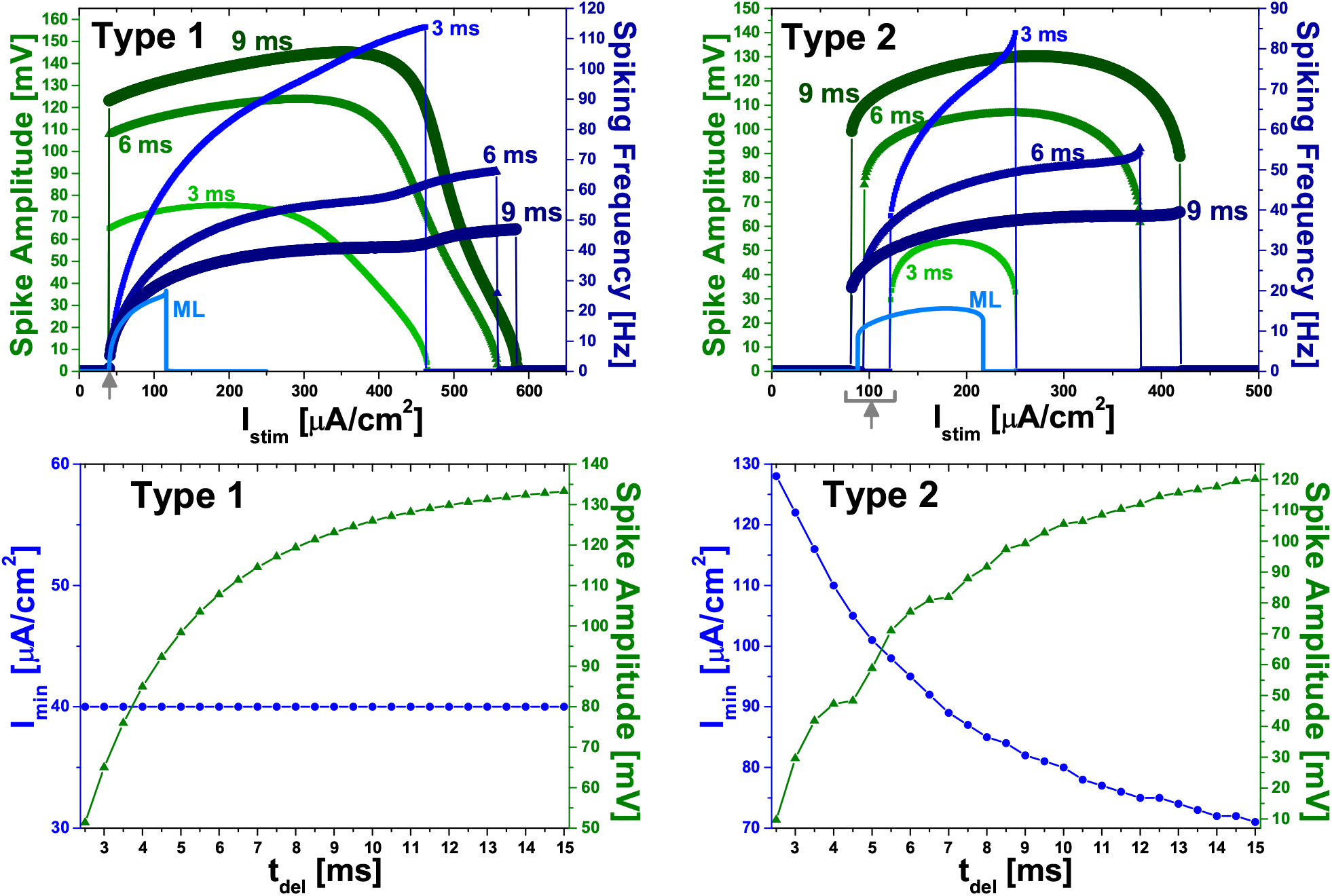
For the SDML model of the 1st (left graphs) and 2nd (right graphs) excitability type, the top graphs show the dependencies of the spike amplitude (left green scale and green curves) and spiking frequency (right blue scale and blue curves) on constant stimulating current *I*_*stim*_ for time delay *t*_*del*_ = 3 ms (thin curves of filled small-size squares), 6 ms (thicker curves of filled midsize triangles), and 9 ms (thickest curves of filled large-size circles). For comparison, the spiking frequency for the standard ML model of the corresponding excitability types is shown by the solid light-blue line. The minimal values of *I*_*stim*_ for generating periodic spikes, denoted further by *I*_*min*_, are marked with the gray upward-pointing arrow. In turn, the bottom graphs show dependencies of the minimal stimulating current *I*_*min*_ (left blue scale and blue curves with filled circles) and the spike amplitude (right green scale and green curves with filled triangles) on time delay *t*_*del*_.

A striking difference between the 1st and 2nd types of excitability can be seen in the corresponding dependencies of *I*_*min*_ on *t*_*del*_: for the 1st type, *I*_*min*_ = 40 *μ*A/cm^2^ and does not depend on *t*_*del*_ in the entire range 2.5 ms *≤ t*_*del*_ *≤* 15 ms, and, for the 2nd type, an explicit hyperbolic-like dependence *I*_*min*_(*t*_*del*_) is observed (see the blue curves on the lower graphs in Fig. 3).

Finally, another striking difference occurs at small values of *t*_*del*_, in particular, at *t*_*del*_ = 1.5 ms and 2 ms. For the ML model parameters resulting in the 2nd excitability type, at these values of *t*_*del*_ periodic generation of spikes in the SDML model does not occur at any value of *I*_*stim*_. In turn, for the parameters leading to the 1st type in the ML model, the SDML model at small *t*_*del*_ values changes its excitability type to the 2nd one (Fig. 4) [23]. Specifically, at *t*_*del*_ = 1.5 ms, periodic spikes appear sharply with high frequency 156 .3 Hz at *I*_*min*_ = 84 *μ*A/cm^2^ that is typical for the 2nd excitability type. At lower values of *I*_*stim*_, periodic spikes are preceded by damped oscillations of the neuron potential, which are also a sign of the 2nd type (e.g., see [24]). At *t*_*del*_ = 2 ms, the picture is similar, although in this case *I*_*min*_ = 40 *μ*A/cm^2^, i.e., it coincides with *I*_*min*_ value for the true 1st excitability type (see the left graphs in Fig. 3). Nevertheless, in this case periodic spikes at *I*_*stim*_ = *I*_*min*_ also occur at a sufticiently high frequency (62.5 Hz) straight away notice a pronounced latency of the occurrence. Thus, these results for the SDML model indicate that i at small values of *t*_*del*_ (≲ 2 ms) only the 2nd excitability type is possible, and ii for the model parameters corresponding to the 1st type in the ML model, the 2nd type transforms into the 1st type with increasing *t*_*del*_ (this transition occurs in the range 2 ms *< t*_*del*_ *<* 2.5 ms).

**Figure 4.**
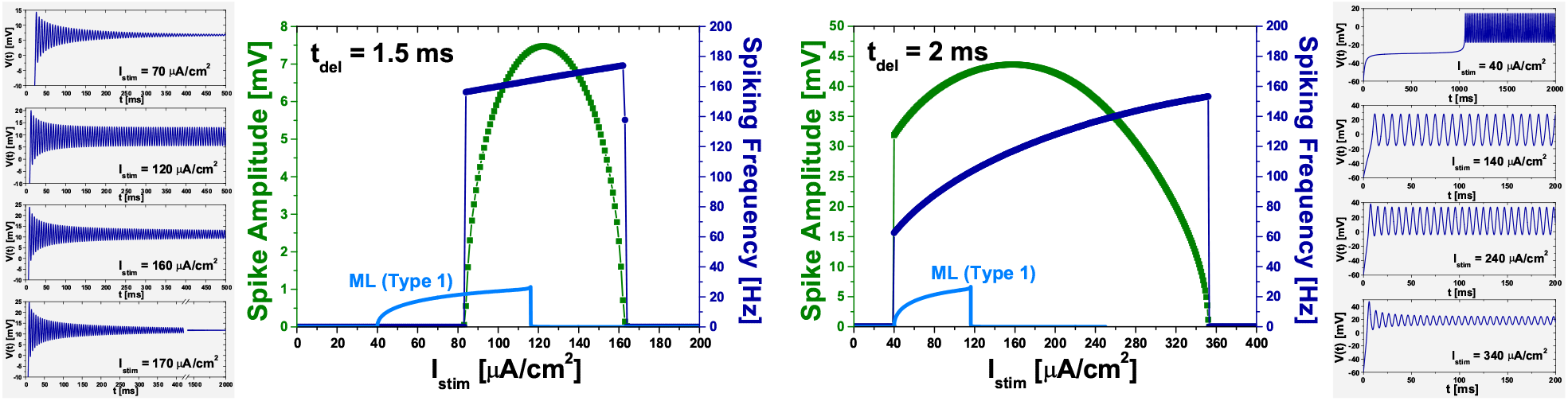
Two central graphs: Dependencies of the spike amplitude (left green scale and green curves) and spiking frequency (right blue scale and blue curves) of the SDML model neuron on constant stimulating current *I*_*stim*_ at time delay *t*_*del*_ = 1.5 ms (left graph) and *t*_*del*_ = 2 ms (right graph), for the model parameters corresponding to the 1st excitability type in the original ML model. The spiking frequency for the latter is shown by the solid light-blue line. In the SDML model, however, the graphs show the discontinuous occurrence of spiking frequency meaning that the excitability type has changed to the 2nd one. The four gray-background graphs on both sides show examples of dynamics of the neuron potential in the SDML model at *t*_*del*_ = 1.5 ms (the leftmost graphs) and *t*_*del*_ = 2 ms (the rightmost graphs) for specific *I*_*stim*_ values indicated on each graph.

## 4. Conclusion

The classical two-dimensional Morris-Lecar (ML) model has been elementary simplified to a single delay differential equation (referred as “Simplified Delay-based Morris-Lecar (SDML) model”), which preserves the occurrence of spikes upon stimulation of the model neuron by direct current. With this simplification, both (i) the initial type of neuronal excitability and (ii) the biophysically realistic waveform of a spike are also preserved in a relatively wide range of the time delay parameter. An additional observation is that a simulation (by the standard Euler method, see Supplementary Material) of the SDML model runs noticeably faster than that of the ML model. It could be beneficial for modeling large neuronal networks, where the ML model is used for single neuron dynamics (cf. [25]).

## Data and code availability

The Supplementary Material for this article contains the data for all graphs in the Figures and ready-to-use MATLAB/Octave codes to reproduce the simulations. It is available online at https://doi.org/10.6084/m9.figshare.18480767.

## Acknowledgments

The author thanks T.Yu. Konovalova for verifying some of the results.

## References

[1] G. E. Hutchinson, Circular causal systems in ecology, Ann. N.Y. Acad. Sci. 50, 221–246 (1948). https://doi.org/10.1111/j.1749-6632.1948.tb39854.x

[2] A. Yu. Kolesov, N. Kh Rozov, The theory of relaxation oscillations for Hutchinson equation, Sb. Math. 202, 829–858 (2011). https://doi.org/10.1070/SM2011v202n06ABEH004168

[3] T. Erneux, Applied Delay Differential Equations (Springer, 2009). https://doi.org/10.1007/978-0-387-74372-1

[4] A.L. Hodgkin, The local electric changes associated with repetitive action in a non-medullated axon, J. Physiol. 107, 165–181 (1948). https://doi.org/10.1113/jphysiol.1948.sp004260

[5] C. Morris, H. Lecar, Voltage oscillations in the barnacle giant muscle fiber, Biophys. J. 35, 193–213 (1981). https://doi.org/10.1016/S0006-3495(81)84782-0

[6] J. Rinzel, B. Ermentrout, Analysis of neural excitability and oscillations, In: C. Koch, I. Segev (Eds.), Methods in Neuronal Modeling: From Synapses to Networks (2nd d., MIT Press, 1998), pp. 251–291. https://www.researchgate.net/publication/237128320

[7] R. FitzHugh, Impulses and physiological states in theoretical models of nerve membrane, Biophys. J. 1, 445–466 (1961). https://doi.org/10.1016/S0006-3495(61)86902-6

[8] J. Nagumo, S. Arimoto, S. Yoshizawa, An active pulse transmission line simulating nerve axon, Proc. IRE 50, 2061–2070 (1962). https://doi.org/10.1109/]RPROC.1962.288235

[9] A. Hodgkin, A. Huxley, A quantitative description of membrane current and its application to conduction and excitation in nerve, J. Physiol. 117, 500 544 (1952). https://dx.doi.org/10.1113/jphysiol.1952.sp004764

[10] V.V. Mayorov, I. Yu. Myshkin, Mathematical modeling of network neurons on the basis of equations with delay [published in Russian], Mat. Model. 2, 64–76 (1990). http://mi.mathnet.ru/eng/mm2481

[11] S.A. Kaschenko, V.V. Mayorov, On a differential-difference equation modeling neuron impulse activity [published in Russian], Mat. Model. 5, 13 25 (1993). http://mi.mathnet.ru/eng/mm2027

[12] S.D. Glyzin, A. Yu. Kolesov, N. h. Rozov, Relaxation self-oscillations in neuron systems: I, Diff. Equat. 47, 927–941 (2011). https://doi.org/10.1134/S0012266111070020

[13] V. Volman et al., Generative modelling of regulated dynamical behavior in cultured neuronal networks, Physica A 335, 249–278 (2004). https://doi.org/10.1016/j.physa.2003.11.015

[14] W. Xiao, H. Gu, M. Liu, Spatiotemporal dynamics in a network composed of neurons with different excitabilities and excitatory coupling, Sci. China Technol. Sci. 59, 1943–1952 (2016). https://doi.org/10.1007/s11431-016-6046-x

[15] A. Calim et al., Chimera states in networks of type-I Morris-Lecar neurons, Phys. Rev. 98, 062217 (2018). https://doi.org/10.1103/PhysRevE.98.062217

[16] K.C.A. Wedgwood et al., Robust spike timing in an excitable cell with delayed feedback, J. R. Soc. Interface 18, 20210029 (2021). https://doi.org/10.1098/rsif.2021.0029

[17] K. Ikeda, J.M. Bekkers, Autapses, Curr. Biol. 16, R308 (2006). https://doi.org/10.1016/j.cub.2006.03.085

[18] C. Wang et al., Formation of autapse connected to neuron and its biological function, Complexity 2017, 5436737 (2017). https://doi.org/10.1155/2017/5436737

[19] R. Swain, The Morris-Lecar Equations with Delay (M.Sc. Thesis, Memorial University of Newfoundland, Canada, 2003). https://research.library.mun.ca/7022/3/Swain_Robin.pdf

[20] X. Song, H. Wang, Y. Chen, Autapse-induced firing patterns transitions in the Morris Lecar neuron model, Nonlinear Dyn. 96, 2341–2350 (2019). https://doi.org/10.1007/s11071-019-04925-7

[21] J.M. Gonzalez-Miranda, Pacemaker dynamics in the full Morris-Lecar model, Commun. Non-linear Sci. Numer. Simulat. 19, 3229–3241 (2014). https://doi.org/10.1016/j.cnsns.2014.02.020

[22] A.V. Paraskevov, T.S. Zemskova, Analytical solution of linearized equations of the Morris-Lecar neuron model at large constant stimulation, Phys. Lett. A 402, 127379 (2021). https://doi.org/10.1016/j.physleta.2021.127379

[23] S.A. Prescott et al., Pyramidal neurons switch from integrators in vitro to resonators under in vivo-like conditions, J. Neurophysiol. 100, 3030–3042 (2008). https://doi.org/10.1152/jn.90634.2008

[24] M.A.D. Roa et al., Scaling law for the transient behavior of type-II neuron models, Phys. Rev. E 75, 021911 (2007). https://doi.org/10.1103/PhysRevE.75.021911

[25] E.M. Izhikevich, Which model to use for cortical spiking neurons?, IEEE Trans. Neural Netw. 15, 1063–1070 (2004). https://doi.org/10.1109/TNN.2004.832719

